# Spatial relationships between white matter degeneration, amyloid load and cortical volume in amnestic mild cognitive impairment

**DOI:** 10.1101/441840

**Authors:** Ileana O. Jelescu, Timothy M. Shepherd, Dmitry S. Novikov, Yu-Shin Ding, Benjamin Ades-Aron, Jacqueline Smith, Thomas Vahle, James S Babb, Kent P. Friedman, Mony J. de Leon, James B. Golomb, James E. Galvin, Els Fieremans

## Abstract

The spatial-temporal relationships between gray and white matter (WM) degeneration during preclinical and early symptomatic Alzheimer’s disease are poorly understood. We characterized β-amyloid deposition, cortical volume and WM degeneration in 44 subjects including healthy control (N=23), amnestic mild cognitive impairment (aMCI) (N=19), and early Alzheimer’s subjects (N=2). Integrated PET-MRI provided simultaneous measurement of ^18^F-Florbetapir uptake in cortical areas, regional brain volumes from structural MRI, and WM tract integrity metrics from diffusion MRI using biophysical modeling.

Across the cohort of healthy control and aMCIs, cortical volumes correlated poorly with β-amyloid deposition in the same area (*p* < 0.05 only in the posterior cingulate and parietal lobe). WM degeneration correlated significantly with both amyloid and volume of connected cortical areas, but more strongly with volume. Diffusion MRI metrics for WM demyelination and/or axonal loss could therefore provide new biomarkers associated with clinical Alzheimer’s conversion. These WM changes may represent sequential propagation of Alzheimer’s neurodegeneration between functionally connected regions, and/or evidence of direct WM injury during the Alzheimer’s pathology cascade.

## 1. Introduction

The neuropathological hallmarks of Alzheimer’s disease are extra-cellular β-amyloid (Aβ) plaques deposition in the cortex (Divry, 1927) and intra-neuronal neurofibrillary tangles caused by abnormal tau protein aggregation (Wood et al., 1986). Other prominent features include hippocampal and cortical atrophy (Convit et al., 1993; de Leon et al., 1989; de Leon et al., 1988; Seab et al., 1988) and white matter tract degeneration (Brun and Englund, 1986; de la Monte, 1989; George et al., 1986). However, there is no consensus on the spatial and temporal relationships of these neurodegenerative changes in individual patients. Their precise contribution to pathologic and clinical disease progression is also not well understood (Archer et al., 2006; Chetelat et al., 2010).

Aβ appears first in neocortical areas, then in the hippocampus and entorhinal cortex, diencephalic structures, and finally the pons and cerebellum (Braak and Braak, 1991; Clark et al., 2011; Thal et al., 2002). Most current models suggest that Aβ deposition may begin one or two decades before presentation of clinical symptoms, while atrophy is temporally more closely associated with the symptomatic stages of amnestic mild cognitive impairment (aMCI) and early Alzheimer’s disease (Jack et al., 2010), although this is challenged by recent data that suggest neurodegeneration can precede amyloid deposition in sporadic AD (Jack Jr and Holtzman, 2013). Furthermore, Aβ deposition generally occurs in a diffuse pattern, whereas Alzheimer’s-related tau pathology and volume loss demonstrate a more sequential, topographic-specific progression (Braak and Braak, 1991). Structural atrophy typically first involves the medial temporal lobe (MTL), but also affects the parietal lobe (PL) (Jacobs et al., 2012) and the posterior cingulate (PC) cortex (Choo et al., 2010).

White matter degeneration in Alzheimer’s disease, as characterized with diffusion tensor imaging (DTI) (Basser et al., 1994), is gathering increasing attention as an important biomarker of neurodegeneration (Kantarci et al., 2014) that may precede structural atrophy (Agosta et al., 2011; Chang et al., 2015; Choo et al., 2010; de la Monte, 1989). The main white matter tracts affected include fornix (Fx), cingulum (CG) and corpus callosum (CC) (Chang et al., 2015; Fieremans et al., 2013; Kantarci et al., 2014; Mielke et al., 2009). Histopathological studies of white matter in the Alzheimer’s brain demonstrate fiber atrophy, patchy rarefaction of fibers, loss of oligodendrocytes, demyelination, and gliosis (de la Monte, 1989; Gouw et al., 2008; Sjobeck et al., 2006).

This study employs integrated PET-MRI to comprehensively characterize spatial correlations between Aβ deposition, structural volume and white matter degeneration, in a pooled cohort of older healthy controls and aMCI subjects, with two additional Alzheimer’s subjects serving as clinical AD-positive controls. Since there can be discordance between pathological and clinical Alzheimer’s disease in individual patients (Negash et al., 2013), the perspective for this study was not to focus on the clinical status of individual subjects, but on the underlying pathology. We carefully examined *in vivo* spatial correlations between different aspects of Alzheimer’s gray and white matter pathology based on simultaneously acquired ^18^F-Florbetapir PET, structural and diffusion MRI.

Few studies so far have examined these three aspects of Alzheimer’s pathology concomitantly (Kantarci et al., 2014; Rieckmann et al., 2016). While data for Aβ, structural volume and DTI are available in the ADNI2 database, in this study we used a more advanced diffusion MRI acquisition than DTI called diffusion kurtosis imaging (DKI), which provides an estimate of the kurtosis of the distribution of water molecule displacements, a measure of microstructural tissue complexity (Jensen et al., 2005). More importantly, DKI data enables us to extract compartment-specific diffusion parameters in the white matter and provide a more specific characterization of WM microstructural changes. Indeed, both DTI and DKI parameters *per se* lack pathological specificity and cannot distinguish, for example, between demyelination, axonal loss or edema. In contrast, the White Matter Tract Integrity (WMTI) two-compartment model is directly derived from the DTI and DKI data, and provides estimates of axonal water fraction (AWF), intra-axonal diffusivity (*D*_a_), and extra-axonal axial and radial diffusivities (*D*_e,||_ and *D*_e,⊥_) for highly aligned WM tracts (Fieremans et al., 2011; Fieremans et al., 2010). In particular, AWF is the ratio of intra-axonal water to total observable tissue (intra-+ extra-axonal) water, while *D*_e,⊥_ is the diffusivity of extra-axonal water molecules perpendicular to the axonal fibers. While the WMTI diffusion model is based on a certain number of assumptions (high fiber alignment, no observable inter-compartment water exchange within white matter and *D*_a_ ≤ *D*_e_,||) that still need definite biological validation, simulations and animal validation studies of parameter specificity have shown that AWF is most sensitive to axonal loss and/or “patchy” demyelination (i.e. areas demyelinated over a few microns are juxtaposed with areas still fully myelinated), while *D*_e,⊥_ is most sensitive to widespread demyelination (Fieremans et al., 2012; Jelescu et al., 2016; Novikov and Fieremans, 2012). Previous studies demonstrated that WMTI metrics across several WM tracts (splenium of the corpus callosum, optic radiations, corticocortical fibers, forceps major, anterior limb of the internal capsule, corona radiata and cingulum) can discriminate well between healthy controls and aMCI, and between aMCI and Alzheimer’s (Fieremans et al., 2013). The WMTI changes observed also suggested increased vulnerability of late-myelinating tracts to Alzheimer’s disease (Benitez et al., 2013). Investigating Aβ deposition, cortical volume and WM integrity at the same time may give insight into how AD pathology propagates through the stages of the disease.

## 2. Methods

### 2.1 Participants

This prospective study was approved by the local Institutional Review Board and informed consent was documented from all 51 participants. Participants were recruited from two sources at NYU: aMCI and Alzheimer’s disease patients were recruited from memory clinics directed by experienced cognitive/behavioral neurologists, while control and aMCI individuals were recruited from NIH-funded studies of aging and dementia. Each participant underwent a comprehensive evaluation including a detailed interview with the participant and a reliable informant, neurological exam and neuropsychological testing. Diagnoses were independently adjudicated by one author (JEG) who was blinded to biomarker status and identifiable personal data. All available clinical data from the electronic health record and research chart was used. The participant’s clinical data was summarized and staged for this study using the Global Deterioration Scale (Reisberg et al., 1988) where GDS 1 or 2 represented normal cognition only distinguished by subjective reporting on interview of life time changes in memory without changes in function at work or home, GDS 3 represented mild cognitive impairment, and GDS 4 represented mild Alzheimer’s disease. During the adjudication process, diagnoses were assigned according to National Institute of Aging – Alzheimer Association criteria for normal controls and presymptomatic individuals (N = 23, 11 males, 68 ± 5 y/o; (Sperling et al., 2011)), aMCI likely due to Alzheimer’s disease (N = 19, nine males, 70 ± 5 y/o; (Albert et al., 2011)), or mild dementia due to Alzheimer’s disease (N = 2, one 78 y/o male and one 69 y/o female; (McKhann et al., 2011)). The first two groups – normal controls and aMCIs – were age-matched. The two Alzheimer’s subjects were included as clinical AD-positive controls in order to see if they fall in the continuity of the biomarker trends established using controls and aMCIs, but were not included in the correlation estimations. As this was a study of relationships between white matter degeneration, amyloid load and cortical volume, six subjects were excluded due to MCI symptoms more consistent with other dementia syndromes, or focal brain abnormality (e.g., due to prior stroke or previous traumatic brain injury) visible on the MRI that likely significantly contributed to their symptoms. One subject was excluded due to degraded PET image quality.

### 2.2 Experimental

Participants underwent simultaneous MRI and PET examination on a 3-Tesla integrated PET-MRI Biograph mMR scanner (Siemens Healthcare, Erlangen, Germany).

#### 2.2.1 MRI

A 1-mm isotropic resolution anatomical MRI image was acquired for cortical and sub-cortical segmentation (3D MP-RAGE: echo time = 2.98 ms / repetition time = 2.3 s / inversion time = 900 ms / flip angle = 9°).

To derive the diffusion tensor, kurtosis tensor and WMTI model parameters, 140 diffusion-weighted images were acquired as follows: 4 *b* = 0 images, *b* = 250 s/mm^2^ – 6 directions, *b* = 1000 s/mm^2^ – 20 directions, *b* = 1500 s/mm^2^ – 20 directions, *b* = 2000 s/mm^2^ – 30 directions, *b* = 2500 s/mm^2^ – 60 directions. Sequence parameters: twice refocused spin-echo single shot echo planar imaging (EPI), echo time = 96 ms / repetition time = 8.2 s / 50 slices / Resolution: 2.5 mm isotropic.

To correct for EPI distortions related to magnetic field inhomogeneity, six additional *b* = 0 images were acquired with reversed phase-encode direction (posterior-to-anterior instead of anterior-to-posterior).

A FLAIR image (FLuid Attenuated Inversion Recovery) was also acquired to evaluate white matter lesion load (32 axial slices / slice thickness = 5 mm / in-plane resolution = 750×692 μm^2^ / echo time = 91 ms / repetition time = 8 s / inversion time = 2.37 s / flip angle = 150°).

#### 2.2.2 PET

For each subject, 9 mCi of ^18^F-Florbetapir (Eli Lilly, Indianapolis, IN, USA) was injected intravenously and 20 minutes of PET list-mode data, starting at 40 minutes post-injection, were reconstructed (Wong et al., 2010) using the Siemens e7tools.

An MRI-based attenuation map was acquired for PET attenuation correction using the VB20 Siemens ultrashort echo time (UTE) sequence (echo time 1 = 0.07 ms / echo time 2 = 2.46 ms / Resolution: 1.6 mm isotropic). The UTE-based attenuation map, which accounts for air, bone and soft-tissue, has shown good concordance with gold-standard CT attenuation maps in the brain, with PET activity underestimation within 3.6% and overestimation within 5.2% depending on the region (Aasheim et al., 2015).

Other PET reconstruction parameters were – algorithm: OP-OSEM (ordinary Poisson ordered subset expectation maximization) with 3 iterations and 21 subsets; matrix: 344×344; 2 mm-kernel Gaussian filter; zoom 2.

### 2.3 Image processing

Figure 1 illustrates the regions of interest (ROIs) considered and their connections.

**Figure 1.**
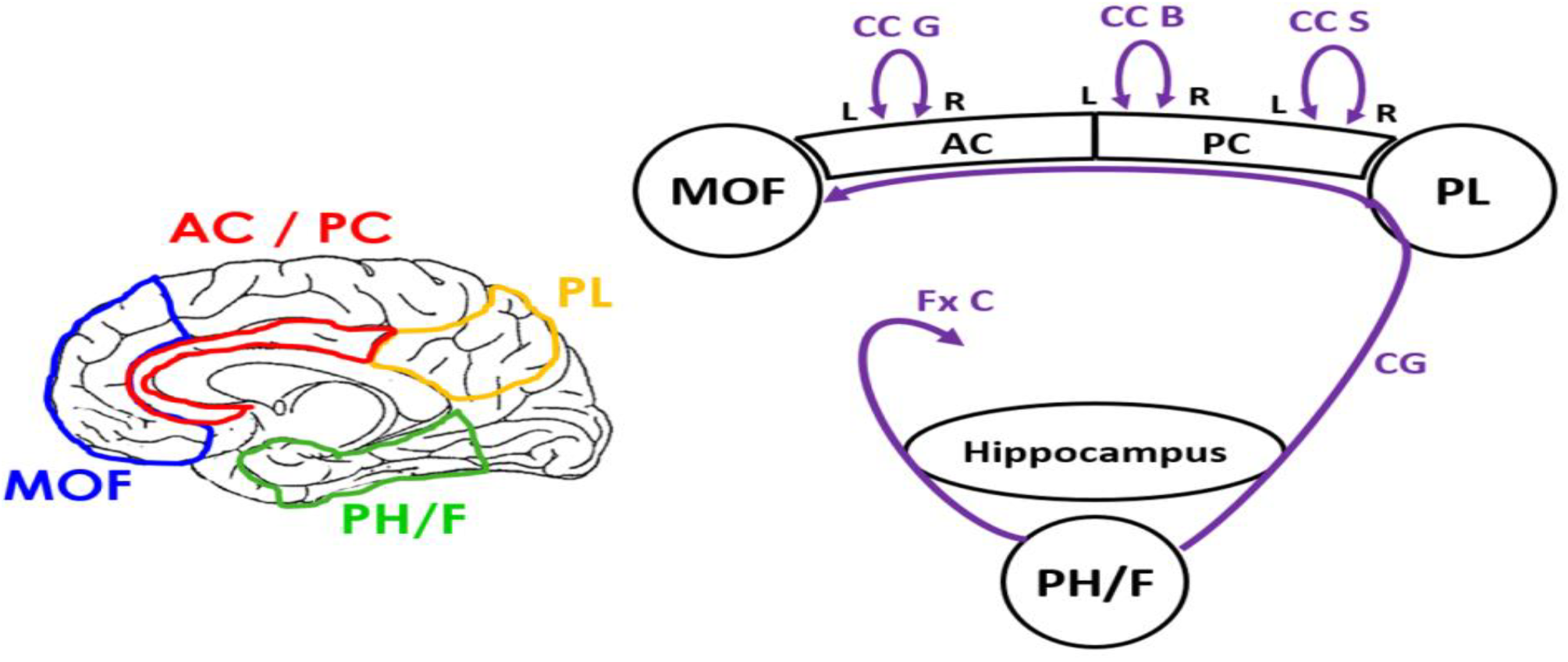
Conceptual model of regions most likely affected by early Alzheimer’s pathology. **Left**: localization of cortical ROIs in the brain. PL: inferior parietal, superior parietal and precuneus; PH/F: entorhinal cortex, fusiform and parahippocampal gyri. **Right**: cortical regions are in black and WM tracts in purple. The corpus callosum interconnects left/right anterior and posterior cingulate. *WM tracts: CC G, CC B and CC S: genu, body and splenium of corpus callosum, respectively; CG: cingulum; Fx C: fornix crus*.

#### 2.3.1 Structure volume

Automated cortical and subcortical segmentation was performed on the high-resolution anatomical T_1_-weighted image using FreeSurfer v5.3.0 (http://surfer.nmr.mgh.harvard.edu/). The volumes of the defined cortical regions of interest – hippocampus alone, parahippocampal gyri, fusiform and entorhinal cortex (PH/F), PL (comprising inferior parietal, superior parietal and precuneus), posterior cingulate cortex (PC), anterior cingulate cortex (AC) and medial orbitofrontal cortex (MOF), see Figure 1 – were normalized to the total intra-cranial volume of each subject.

#### 2.3.2 Florbetapir uptake

The reconstructed PET image was registered to the MRI anatomical image using FSL’s linear registration “FLIRT” (Jenkinson et al., 2002) with a mutual information cost function. Using the structural masks obtained from FreeSurfer, the mean standardized uptake value (SUV) was calculated in each of the cortical regions known for pathological uptake of florbetapir (AC, PC, MOF, PL and PH/F), and was further normalized to the mean SUV in the whole cerebellum to yield relative SUV (SUVr) (Clark et al., 2011).

#### 2.3.3 Diffusion metrics

The diffusion-weighted images were corrected for motion, eddy current and field inhomogeneity distortions using FSL’s “Topup” and “EDDY”. The diffusion and kurtosis tensors were estimated in each voxel of the brain using a weighted linear least squares algorithm (Veraart et al., 2013) from which parametric maps of relevant DKI (FA, MD, MK) and WMTI (AWF, *D*_e,⊥_) metrics were calculated. Each subject’s FA map was registered to the Johns Hopkins University FA template (Oishi et al., 2009) using non-rigid registration (FSL’s “FNIRT” (Andersson et al., 2010)) and the white matter tracts of interest (corpus callosum genu (CCG), body (CCB) and splenium (CCS), CG – encompassing both cingulate cingulum and parahippocampal cingulum – and FxC, see Figure 1) were imported from the JHU atlas back into subject space. For DKI metrics, the entire white matter ROIs were considered in the analysis; for WMTI, an additional FA threshold of 0.4 was applied on the JHU template to ensure selection of voxels with highly unidirectional fiber orientation necessary for the model applicability. Mean values of DKI and WMTI metrics were calculated for each white matter ROI.

#### 2.3.4 White matter lesion load

White matter lesions were segmented on the FLAIR images using *FireVoxel*, a semi-automatic integrated imaging data processing software, and reviewed by a board-certified neuroradiologist. The total white matter lesion volume was subsequently calculated and normalized to the total intra-cranial volume.

### 2.4 Correlations

The focus of this study was on early Alzheimer’s pathology rather than clinical status. We therefore pooled and analyzed data from the entire cohort of control and aMCI subjects, who were likely to have different degrees of underlying Alzheimer’s pathology. The impact of age was regressed out in all analyses. In a separate analysis, both age and WM lesion load were used as covariates in the correlation estimations.

The following partial Pearson correlations, co-varying for either subject age or both subject age and WM lesion load, were calculated and significance level was set to *p* < 0.05:

– between the SUVr’s in the pre-defined cortical areas (see Figure 1) to assess whether Aβ is generalized, focal or topographically sequential. A Bonferroni correction was further applied to account for correlations to four different ROIs (*p* < 0.0125);
– between the structure volumes in the pre-defined cortical areas and hippocampus (see Figure 1) to assess whether atrophy is generalized, focal or topographically sequential. A Bonferroni correction was further applied to account for correlations to five different ROIs (*p* < 0.01);
– between the SUVr and the volume of each cortical ROI (MOF, AC, PC, PL and PH/F) to assess whether Aβ deposition and structural volume are related;
– between the volume of a cortical ROI and the diffusion metrics in connected white matter tracts (see Figure 1) to assess whether structural volume and white matter degeneration are related. A Bonferroni correction was further applied to account for correlations to five different diffusion metrics (*p* < 0.01);
– between the SUVr in a cortical ROI and the diffusion metrics in connected white matter tracts (see Figure 1) to assess whether Aβ deposition and white matter degeneration are related. For these correlations, corresponding cortical volume was also used as a covariate together with age, in order to test whether WM degeneration and cortical SUVr are directly or indirectly correlated. A Bonferroni correction was further applied to account for correlations to five different diffusion metrics (*p* < 0.01).

## 3. Results

Cortical SUVr values were 1.04 ± 0.10 in controls, 1.13 ± 0.15 in aMCI and 1.50 ± 0.03 in the two AD subjects. These values were fully consistent with the ranges previously reported for healthy controls (1.05 ± 0.16), aMCI (1.20 ± 0.28) and Alzheimer’s patients (1.40 ± 0.27) (Johnson et al., 2013) obtained after manual segmentation on the PET images (without an MRI reference) but with otherwise identical methods. All aMCI subjects were amyloid positive (mean SUVr > 1.1 (Joshi et al., 2012)) in at least two lateralized cortical regions, while the two AD subjects were amyloid positive in all cortical areas examined. The range of hippocampal volumes was 0.16 – 0.38 (% total intracranial volume), with 0.26 ± 0.04 in controls, 0.24 ± 0.05 in aMCIs and 0.23 ± 0.03 in AD. These ranges were also in agreement with the literature on healthy controls (0.34 ± 0.06), aMCI (0.29 ± 0.07) and Alzheimer’s patients (0.25 ± 0.06) obtained from segmentation of structural MRI images (He et al., 2012). Our subject pool therefore contained the range of typically observed metrics for healthy control, aMCI and AD.

The correlations obtained when regressing out the age only, or both age and WM lesion load were very similar: correlation coefficients varied by 0.03 (in absolute value) at most, with no consistent trend of weakened or strengthened correlations. Only correlations with subject age as a covariate are therefore reported and discussed in the following sections.

### 3.1 Cortical correlations for regional brain volume and Aβ deposition

*Figure 2A* illustrates statistically significant correlations between the SUVr’s of cortical areas that are directly connected. All correlation coefficients *r* and *p*-values are listed in *Table 1*. The SUVr’s of all cortical regions were highly correlated, but the strongest correlations were between ROIs that we model as being directly connected by a WM tract. The strongest correlation was in the frontal region, between the SUVr’s in MOF and AntCing (*r* = 0.97, *p* < 0.001) (*Figure 3*). The smallest observed correlation was between the SUVr’s in the AC and PH/F (*r* = 0.78, *p* < 0.001). These results are consistent with established widespread patterns of Aβ deposition in Alzheimer’s disease (Braak and Braak, 1991).

**Figure 2.**
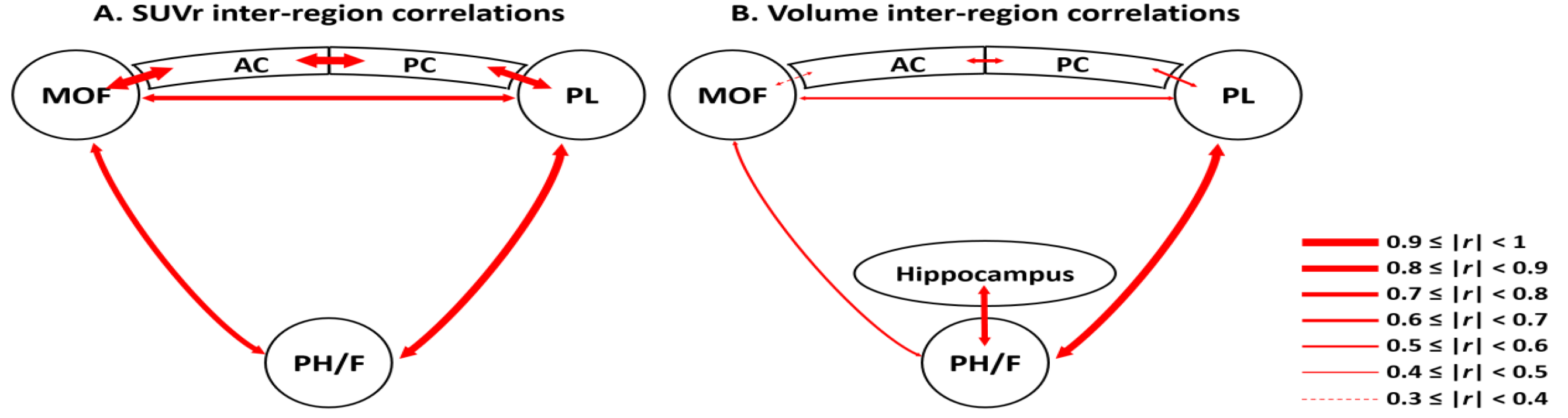
Significant correlations between SUVr’s (A) or volumes (B) of cortical regions directly connected by WM tracts. The thickness of the red arrows illustrates the strength of the correlation. All correlation values are provided in Tables 1 and 2.

**Figure 3.**
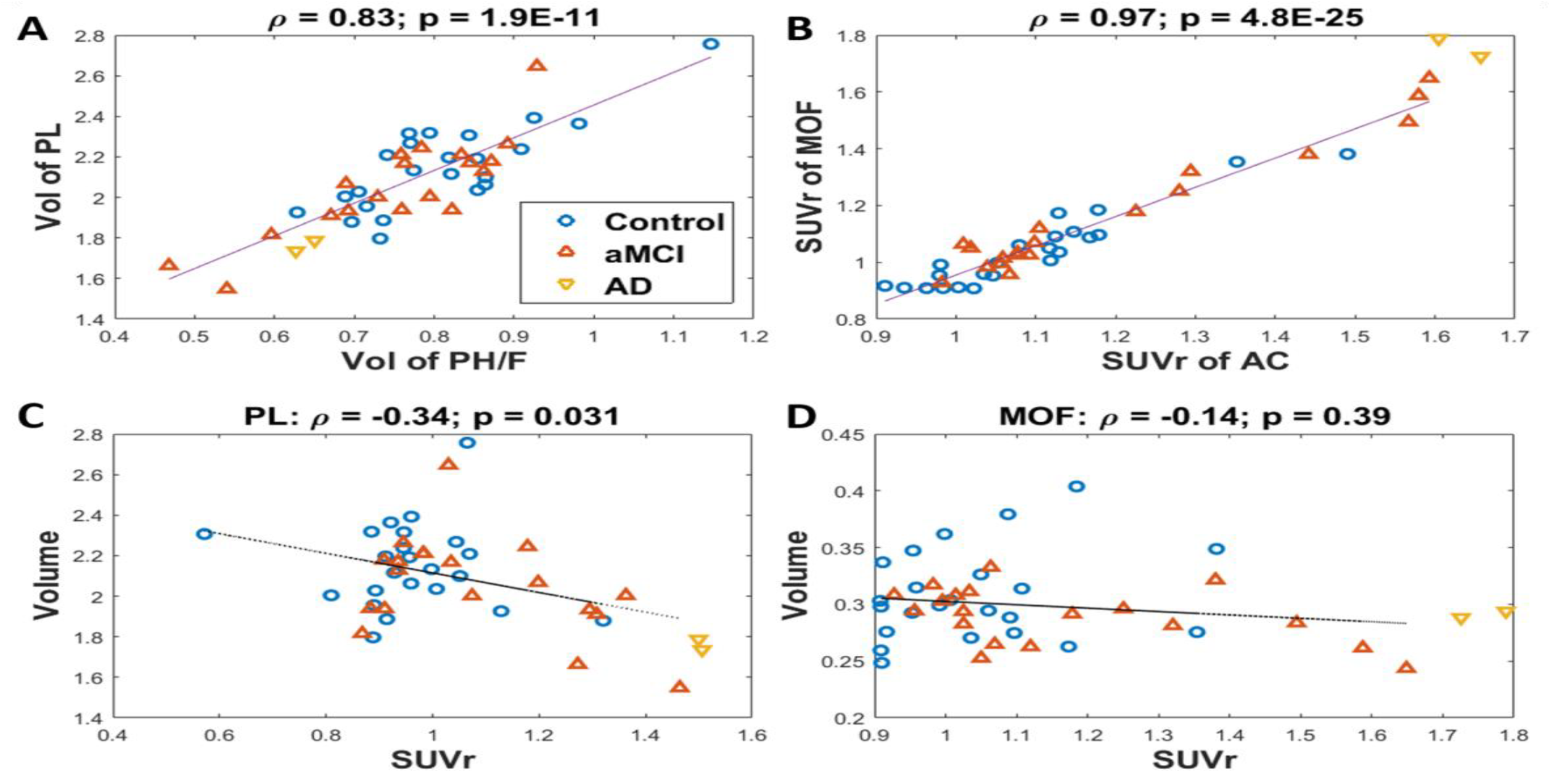
Examples of correlations between SUVr and volume (listed in Table 3). A: The volumes of PH/F and of PL correlate most strongly with each other. B: The SUVr’s of AC and MOF correlate most strongly with each other. The strongest correlation between volume and SUVr in a given cortical area is in the PL (C), while the weakest is in the MOF (non-significant) (D). Note the Pearson ρ and p-value account for age as a covariate, while the solid black line illustrates a direct linear correlation (no covariates). All correlation values are visualized in Figure 2 and listed in Tables 1 and 2.

**Table 1.** Partial Pearson correlations, co-varying for subjects’ age, between SUVr’s in cortical areas. r: correlation coefficient; p: p-value. Green: Significant correlations after Bonferroni correction, each ROI being correlated to four others (p < 0.0125).

| SUVR | PH/F |  | PL |  | PC |  | AC |  | MOF |  |
| --- | --- | --- | --- | --- | --- | --- | --- | --- | --- | --- |
|  | *r* | *p* | *r* | *p* | *r* | *p* | *r* | *p* | *r* | *p* |
| PH/F |  |  | 0.87 | 1.06E-13 | 0.80 | 5.28E-10 | 0.78 | 2.63E-09 | 0.79 | 9.62E-10 |
| PL | 0.87 | 1.06E-13 |  |  | 0.88 | 2.25E-14 | 0.82 | 8.58E-11 | 0.79 | 6.32E-10 |
| PC | 0.80 | 5.28E-10 | 0.88 | 2.25E-14 |  |  | 0.94 | 1.81E-20 | 0.90 | 1.21E-15 |
| AC | 0.78 | 2.63E-09 | 0.82 | 8.58E-11 | 0.94 | 1.81E-20 |  |  | 0.97 | 4.84E-25 |
| MOF | 0.79 | 9.62E-10 | 0.79 | 6.32E-10 | 0.90 | 1.21E-15 | 0.97 | 4.84E-25 |  |  |

*Figure 2B* illustrates statistically significant correlations between volumes for GM regions that are directly connected. See *Table 2* for all correlation coefficients *r* and *p*-values. All correlations evaluated were significant, but, unlike florbetapir SUVr, the strength of the correlations varied greatly between different regions. The strongest correlations were between PH/F and PL volumes (*r* = 0.83, *p* < 0.001) (*Figure 3*), and between hippocampal and PL volumes (*r* = 0.78, *p* < 0.001), while the weakest was between MOF and AC (*r* = 0.34, *p* < 0.05). These results illustrate that atrophy in early Alzheimer’s pathology is more tightly correlated to the hippocampus, PH/F and PL.

**Table 2.** Partial Pearson correlations, co-varying for subjects’ age, between volumes of cortical areas and hippocampus. r: correlation coefficient; p: p-value. Orange: significant correlations (p < 0.05); Green: Significant correlations after Bonferroni correction, each ROI being correlated to five others (p < 0.01).

| Volume | PH/F |  | PL |  | PC |  | AC |  | MOF |  | Hipp. |  |
| --- | --- | --- | --- | --- | --- | --- | --- | --- | --- | --- | --- | --- |
|  | *r* | *p* | *r* | *p* | *r* | *p* | *r* | *p* | *r* | *p* | *r* | *p* |
| PH/F |  |  | 0.83 | 1.94E-11 | 0.57 | 8.58E-05 | 0.49 | 1.24E-03 | 0.57 | 1.07E-04 | 0.76 | 7.67E-09 |
| PL | 0.83 | 1.94E-11 |  |  | 0.60 | 3.52E-05 | 0.50 | 8.06E-04 | 0.53 | 3.78E-04 | 0.78 | 1.27E-09 |
| PC | 0.57 | 8.58E-05 | 0.60 | 3.52E-05 |  |  | 0.62 | 1.30E-05 | 0.42 | 6.38E-03 | 0.57 | 9.85E-05 |
| AC | 0.49 | 1.24E-03 | 0.50 | 8.06E-04 | 0.62 | 1.30E-05 |  |  | 0.34 | 3.10E-02 | 0.49 | 1.02E-03 |
| MOF | 0.57 | 1.07E-04 | 0.53 | 3.78E-04 | 0.42 | 6.38E-03 | 0.34 | 3.10E-02 |  |  | 0.62 | 1.30E-05 |
| Hipp. | 0.76 | 7.67E-09 | 0.78 | 1.27E-09 | 0.57 | 9.85E-05 | 0.49 | 1.02E-03 | 0.62 | 1.30E-05 |  |  |

More importantly, correlations between SUVr and volume in a given cortical area were very limited and were only significant in the PL (*r* = −0.34, *p* < 0.05) and in the PC (*r* = −0.32, *p* < 0.05) (*Table 3*).

**Table 3.**
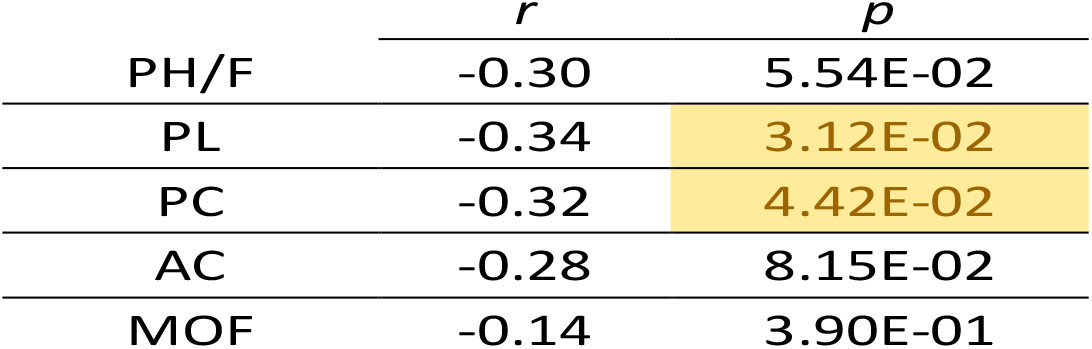
Correlations between SUVr and volume in cortical areas. Yellow: significant correlations.

### 3.2 White matter pathology correlations to regional brain volume and Aβ deposition

Overall there were strong correlations between cortical volumes and WM diffusion parameters (Figure 4A-B and Supplementary Table 1). Significant correlations between cortical florbetapir SUVr and WM diffusion also were found, predominantly in MOF, AC and PC (Figure 4C-D, *Supplementary Table 2*).

**Figure 4.**
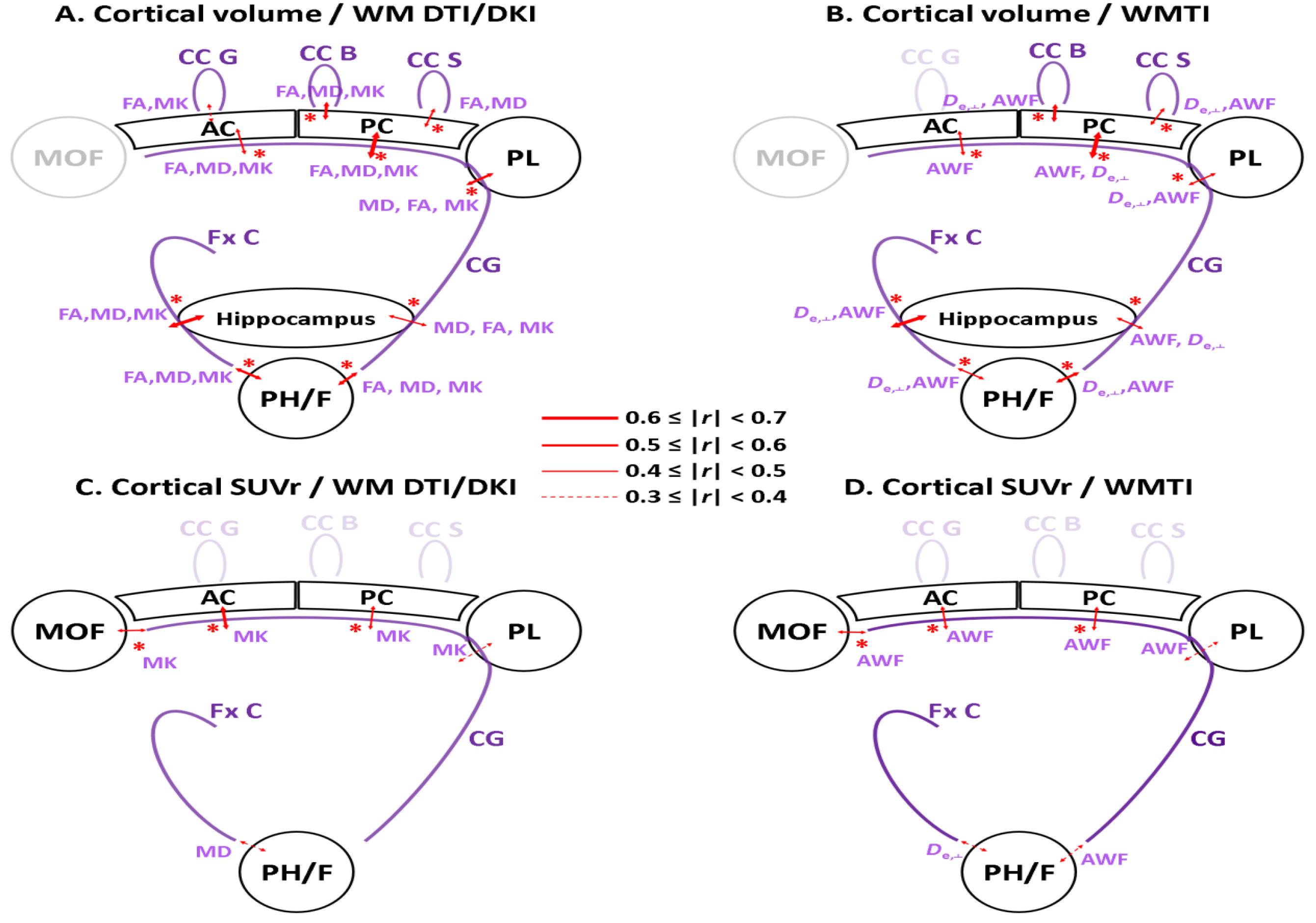
Significant correlations between diffusion metrics in the WM tracts and the volume of related cortical regions (A-B), or the SUVr of related cortical regions (C-D). All diffusion metrics with a significant correlation are listed in purple at the corresponding location. The thickness of the red arrow is based on the strongest correlation among all DTI/DKI or WMTI metrics, respectively. The red asterisks indicate correlations that remain significant after Bonferroni correction (*p* < 0.01). Cortical regions or WM tracts without significant correlation are whited-out for clarity. All correlation values are provided in Supplementary Tables 1 and 2.

All cortical ROIs examined, with the exception of the MOF, displayed correlations between their volume and diffusion metrics in connected WM tracts (Supplementary Table 1). The strongest correlations are plotted in *Figure 5A-B*. In the crus of the fornix and the cingulum, all three tensor metrics considered (FA, MD, MK) correlated strongly with the volume of connected cortical areas (hippocampus and PH/F for the fornix, and hippocampus, PH/F, PL, PC and AC for the cingulum). Diffusion tensor metrics in the corpus callosum also correlated with anterior and posterior cingulate volumes. Lower cortical volumes were associated with lower white matter FA and MK, and with higher MD, i.e. cortical damage correlated with WM damage.

**Figure 5.**
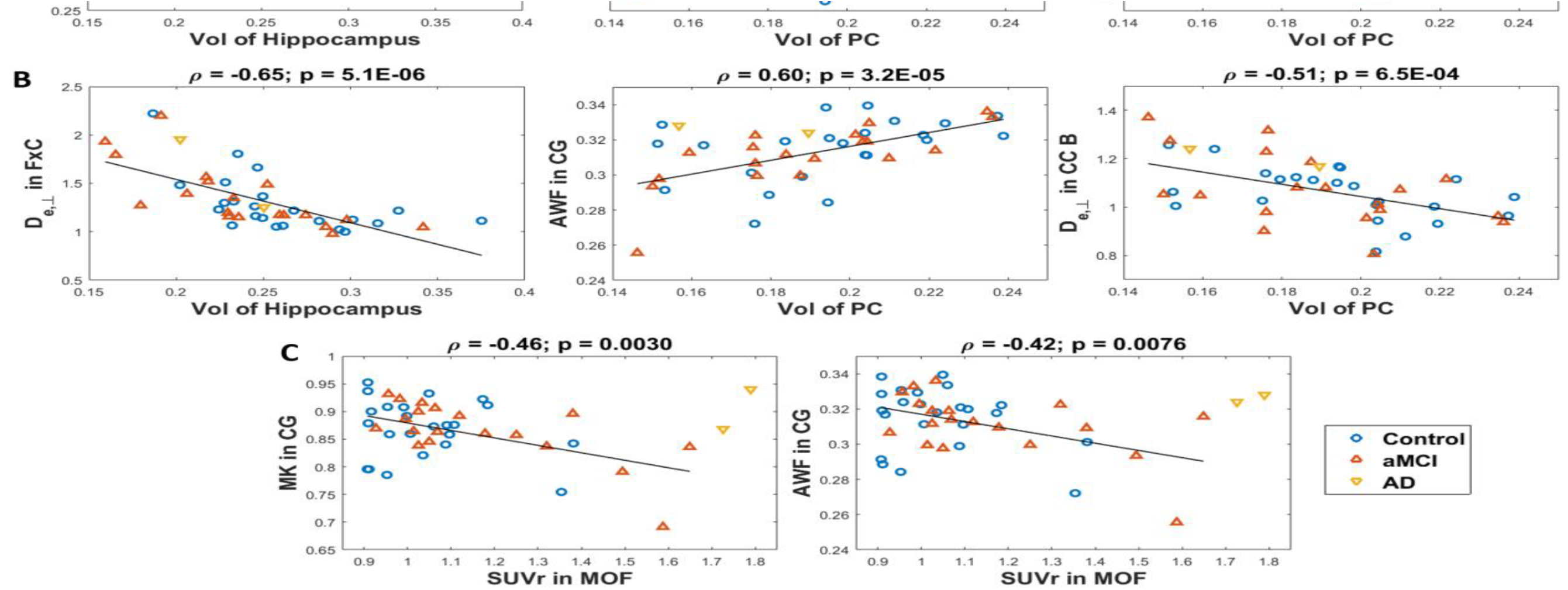
Strongest correlations between volumes of cortical regions and diffusion metrics in connecting WM tracts, in terms of DTI/DKI (A) and WMTI (B). Note the Pearson ρ and p-value account for age as a covariate, while the solid black line illustrates a direct linear correlation (no covariates). Using both age and cortical volume as a covariate, significant correlations remain between SUVr in the MOF and diffusion metrics in the CG (C). All correlation values are visualized in Figure 4 and listed in Supplementary Tables 1 and 2.

The strongest correlations involving DTI and DKI metrics also translated into significant correlations for the WMTI metrics. Lower cortical volumes were associated with lower white matter AWF (i.e. axonal loss or patchy demyelination) and higher *D*_e,⊥_ (i.e. uniform demyelination). In the FxC, CC body and splenium, correlations were significant in terms of both AWF and *D*_e,⊥_, with the latter being stronger. In the cingulum, it was AWF mostly that correlated with volume of the AC, PC and hippocampus.

Nearly all correlations remained significant after Bonferroni correction for multiple comparisons (5 diffusion metrics: *p* < 0.01), with the exception of AC volume vs. diffusion metrics in the CC genu.

Diffusion metrics correlated significantly with cortical SUVr’s only in the FxC and CG but suggested that higher cortical amyloid levels were associated with WM damage in connected tracts (see *Supplementary Table 2* for numerical values). Remarkably, the correlations between MK and AWF in the CG and the SUVr’s of MOF, PC and AC remained significant after Bonferroni correction and also after the volume of the corresponding cortical region was regressed out (see *Figure 5C* and *Supplementary Table 3*).

## 4. Discussion

In this work, we focused on three aspects of mild cognitive impairment preceding Alzheimer’s disease and characterized the spatial correlations of Aβ deposition, structural volume and white matter degeneration. We first discuss the study design, then cortical correlations (Aβ burden and structural volume) and finally correlations of WM damage to cortical measures of Aβ burden and volume. We also explore potential limitations of the current study.

The study was carried out in a cohort of older healthy controls and aMCI subjects. Clinical status however was purposefully not used in the analysis because there often can be discordance between *in vivo* biomarkers and cognitive scores due to multiple factors, including cognitive reserve (Negash et al., 2013). The expected range of amyloid deposition and neurodegeneration was confirmed by the range of values in cortical SUVr and structural volumes (*Figure 3* and *Figure 5*), which matched prior findings across this spectrum of clinical disease (He et al., 2012; Johnson et al., 2013). Remarkably, clinical status is not associated with a clear-cut threshold in quantitative biomarker values, with the exception perhaps of SUVr values in AD which are clearly higher than the rest of the cohort’s. Nonetheless, the “healthy” and “diseased” ends of the spectra are typically dominated by control and aMCI/AD subjects, respectively, and all aMCI subjects were found to be amyloid positive in at least two lateralized cortical regions.

Potential confounds were carefully minimized in the study design. The simultaneous PET-MRI acquisition removes confounds of both misregistration and disease progression between the time points of MRI and PET data collection. The effect of normal aging was corrected for by using the subjects’ age as a covariate in all correlation estimates. Additionally, the WM tract segmentation was based on automatic registration to the JHU atlas and not on the individual subject’s FA maps, thus individual WM disease should not confound the segmentation.

### SUVr and structural volume

SUVr inter-region correlations revealed widespread accumulation of Aβ in the cortical areas examined, consistent with previous literature (Braak and Braak, 1991). In contrast, inter-region correlations of structural volume were more localized and strongest between the hippocampus, PH/F and PL, which are known to be early sites of both structural atrophy and tau-pathology (Bobinski et al., 1997; Bobinski et al., 1996; Braak and Braak, 1991; Jack et al., 1997; Jacobs et al., 2012). While this study was cross-sectional in nature, the pattern of correlations between GM regions observed also supports that atrophy may follow a topographic sequential progression, as previously reported (Convit et al., 1997).

There were weak correlations between volume and SUVr in the PL and PC only. Cortical atrophy and Aβ accumulation thus appeared independent for most brain regions, although whole brain atrophy has been shown to correlate with global Aβ uptake (Archer et al., 2006), suggesting the presence of Aβ may still be a requirement to initiate aspects of neurodegeneration and the eventual development of clinical Alzheimer’s disease in most patients. Our results are concordant with previous studies that suggested spatial and temporal dissociations between amyloid status and atrophy (Huijbers et al., 2015; Insel et al., 2015; Jack et al., 2014; Wang et al., 2015). Furthermore, atrophy is likely a relatively late manifestation of underlying Alzheimer’s pathology – other cortical pathological changes such as synaptic loss that are not easily observed with current PET-MRI techniques may occur following Aβ deposition prior to cortical thinning and volume loss (Scheff and Price, 2003).

### WM degeneration

In this cross-sectional study, diffusion biomarkers of degeneration in the examined white matter tracts correlated strongly with the structural volume of cortical regions connected by those tracts, similar to previous DTI studies (Choo et al., 2010; Kantarci et al., 2014), except in the frontal areas (MOF volume did not correlate with diffusion metrics in the CG). This result suggests regional brain volume loss and white matter pathology are already tightly linked early on in the disease.

Remarkably, in the frontal areas (MOF and AC), where correlations between SUVr and volume were absent, and correlations between volume and diffusion in the CG were either weak (AC) or absent (MOF), cortical florbetapir SUVr correlated strongly with diffusion metrics in the cingulum.

Therefore, white matter pathology could either occur closely following cortical atrophy (Braak and Braak, 1991; Zhou et al., 2010), precede cortical atrophy (Chang et al., 2015; Choo et al., 2010; de la Monte, 1989), or be an independent process from cortical atrophy (Bartzokis, 2011). However, while WM changes may precede atrophy as MRI-observable phenomena, they could still follow subtle cortical pathology – such as tau inclusions leading to synaptic loss – that could not be evaluated in the current study. A longitudinal study, ideally including markers of tau pathology, may therefore give additional insights into the exact temporal progression of Alzheimer’s disease pathological changes in individual subjects.

Our findings confirm previous DTI studies suggesting that WM damage may play a central role in how Alzheimer’s disease develops and progresses. Here, the characterization of WM damage may be improved by using the WMTI metrics (instead of standard DTI metrics), which are by design more specific for the underlying pathology. Overall, the strong involvement of *D*_e,⊥_ (correlating stronger with volume than AWF) suggests that homogeneous demyelination may affect the FxC and CC body and splenium, while early (patchy) demyelination or axonal loss may be occurring primarily in the cingulum bundle (evidenced by the AWF mostly correlating with volume of the AC, PC and hippocampus). Since WMTI metrics potentially improve specificity, one may expect a trade-off in sensitivity compared to DTI metrics (i.e. in terms of correlation coefficient and *p*-value). However, the loss in sensitivity reported here was minimal and in a few cases even absent (with *D*_e,⊥_ sometimes displaying a higher correlation coefficient than DTI or DKI metrics).

### Limitations and Future Work

This study has some limitations that may be addressed in future. The CG ROI encompasses both cingulate cingulum and parahippocampal cingulum, future analysis of these separately could possible shed more light on the relationships investigated here. Due to the cross-sectional design, we used correlational data to evaluate disease propagation. A future longitudinal study is needed to examine the spatiotemporal progression of these pathological changes in both gray matter and white matter, as well as to elucidate whether in white matter AWF or *D*_e,⊥_ changes occur before, concurrently or after cortical volume loss and Aβ accumulation for particular regions in individual brains. Another potential limitation of the study is that we did not stratify between clinical groups. Although the focus was on quantitative biomarkers, not clinical diagnosis, it would be of interest in a future study to also compare correlations between each of these biomarkers and cognitive performance (e.g. WMTI parameters may provide a structural MRI biomarker for cognitive reserve (Benitez et al., 2014)). While the current study is underpowered to examine the interactions of amyloid burden, cortical volume and WM integrity within clinical groups, and whether these interactions change between groups, larger cohorts should make this evaluation possible in the future.

## 5. Conclusions

We provided comprehensive characterization of spatial correlations between cortical Aβ accumulation, cortical volume and WM degeneration in a cohort of control and aMCI subjects using integrated PET-MRI. The characterization of WM degeneration was performed using metrics with improved specificity compared to DTI – AWF and *D*_e,⊥_, which may provide new biomarkers for Alzheimer’s disease around the time of clinical conversion and improve our understanding of the early propagation of Alzheimer’s pathology. Results suggest that Aβ deposition and structural volume are nearly uncorrelated in any given cortical region. WM degeneration in the form of demyelination and/or axonal loss however correlates very strongly to cortical volume, and remarkably, also correlates to Aβ deposition in frontal areas that are yet spared from detectable atrophy. This places WM degeneration as a potential temporal link between typical early Aβ accumulation and late cortical atrophy. Neurodegeneration, which may depend on tau pathology first observed in the MTL, could propagate to other cortical areas via degeneration of the WM tracts. The spatiotemporal correlations of these biomarkers in individual subjects will be the focus of future work.

## Acknowledgments

The authors thank Emma Ben-Avi, Sonja Blum, MD, Tracy Butler, MD, Stephanie Chrisphonte, MD, Patrick Harvey, Thet Oo MBBS, Matthew Lustberg, Martin Sadowski, MD, PhD, Alok Vedvyas, and Thomas M Wisniewski, MD for help with subject recruitment, and Kimberly Jackson for technical assistance with data acquisition. The radiotracer Amyvid (florbetapir) was provided to the study by Avid Radiopharmaceuticals Inc., Philadelphia, PA, USA. This work was supported by the Alzheimer Drug Discovery Foundation (E.F.), NIH NIA 1K23AG048622-01 (T.M.S), NIH NINDS R01NS088040 (E.F. and D.S.N.), NIH NIA R01AG040211 (J.E.G.), and NIH NIA AG022374, AG13616, AG12101, AG08051 (M.J.dL.). This work was also supported in part by the Center for Advanced Imaging Innovation and Research, a NIH NIBIB Biomedical Technology Resource Center (P41EB017183) and also benefitted greatly from the Alzheimer’s disease center (ADC) program grant (P30AG0851).

## Supplementary Material

**Supplementary Table 1.** Partial Pearson correlations, co-varying for subjects’ age, between WM diffusion metrics and volume of related cortical regions. *r*: correlation coefficient; *p*: *p*-value. Yellow: significant correlations (*p* < 0.05); Green: still significant after Bonferroni correction, accounting for five diffusion metrics (*p* < 0.01).

|  |  | Hippocampus<br>Volume |  | PH/F<br>Volume |  | PL<br>Volume |  | PC<br>Volume |  | AC<br>Volume |  | MOF<br>Volume |  |
| --- | --- | --- | --- | --- | --- | --- | --- | --- | --- | --- | --- | --- | --- |
|  |  | *r* | *p* | *r* | *p* | *r* | *p* | *r* | *p* | *r* | *p* | *r* | *p* |
| FxC | FA | 0.63 | 1.10E-05 | 0.43 | 4.54E-03 |  |  |  |  |  |  |  |  |
|  | MD | -0.61 | 2.14E-05 | -0.56 | 1.41E-04 |  |  |  |  |  |  |  |  |
|  | MK | 0.50 | 8.20E-04 | 0.34 | 2.98E-02 |  |  |  |  |  |  |  |  |
|  | AWF | 0.52 | 4.96E-04 | 0.46 | 2.34E-03 |  |  |  |  |  |  |  |  |
|  | De,⊥ | -0.65 | 5.06E-06 | -0.50 | 8.86E-04 |  |  |  |  |  |  |  |  |
| CG | FA | 0.44 | 3.72E-03 | 0.55 | 2.20E-04 | 0.34 | 2.90E-02 | 0.63 | 8.88E-06 | 0.44 | 3.99E-03 | 0.07 | 6.42E-01 |
|  | MD | -0.48 | 1.34E-03 | -0.52 | 5.41E-04 | -0.55 | 2.19E-04 | -0.43 | 4.71E-03 | -0.31 | 4.54E-02 | -0.15 | 3.55E-01 |
|  | MK | 0.33 | 3.59E-02 | 0.43 | 4.73E-03 | 0.34 | 2.92E-02 | 0.42 | 6.17E-03 | 0.37 | 1.75E-02 | 0.14 | 3.98E-01 |
|  | AWF | 0.45 | 3.52E-03 | 0.48 | 1.42E-03 | 0.43 | 4.91E-03 | 0.60 | 3.15E-05 | 0.44 | 3.84E-03 | 0.05 | 7.40E-01 |
|  | De,⊥ | -0.44 | 4.42E-03 | -0.55 | 2.18E-04 | -0.46 | 2.60E-03 | -0.47 | 2.12E-03 | -0.30 | 5.73E-02 | -0.11 | 5.03E-01 |
| CCS | FA |  |  |  |  |  |  | 0.41 | 8.45E-03 |  |  |  |  |
|  | MD |  |  |  |  |  |  | -0.31 | 4.63E-02 |  |  |  |  |
|  | MK |  |  |  |  |  |  | 0.09 | 5.58E-01 |  |  |  |  |
|  | AWF |  |  |  |  |  |  | 0.32 | 4.34E-02 |  |  |  |  |
|  | De,⊥ |  |  |  |  |  |  | -0.41 | 8.10E-03 |  |  |  |  |
| CCB | FA |  |  |  |  |  |  | 0.50 | 9.10E-04 |  |  |  |  |
|  | MD |  |  |  |  |  |  | -0.50 | 8.23E-04 |  |  |  |  |
|  | MK |  |  |  |  |  |  | 0.46 | 2.34E-03 |  |  |  |  |
|  | AWF |  |  |  |  |  |  | 0.47 | 1.75E-03 |  |  |  |  |
|  | De,⊥ |  |  |  |  |  |  | -0.51 | 6.49E-04 |  |  |  |  |
| CCG | FA |  |  |  |  |  |  |  |  | 0.35 | 2.41E-02 |  |  |
|  | MD |  |  |  |  |  |  |  |  | -0.20 | 2.00E-01 |  |  |
|  | MK |  |  |  |  |  |  |  |  | 0.35 | 2.56E-02 |  |  |
|  | AWF |  |  |  |  |  |  |  |  | 0.31 | 5.23E-02 |  |  |
|  | De,⊥ |  |  |  |  |  |  |  |  | -0.24 | 1.26E-01 |  |  |

**Supplementary Table 2.**
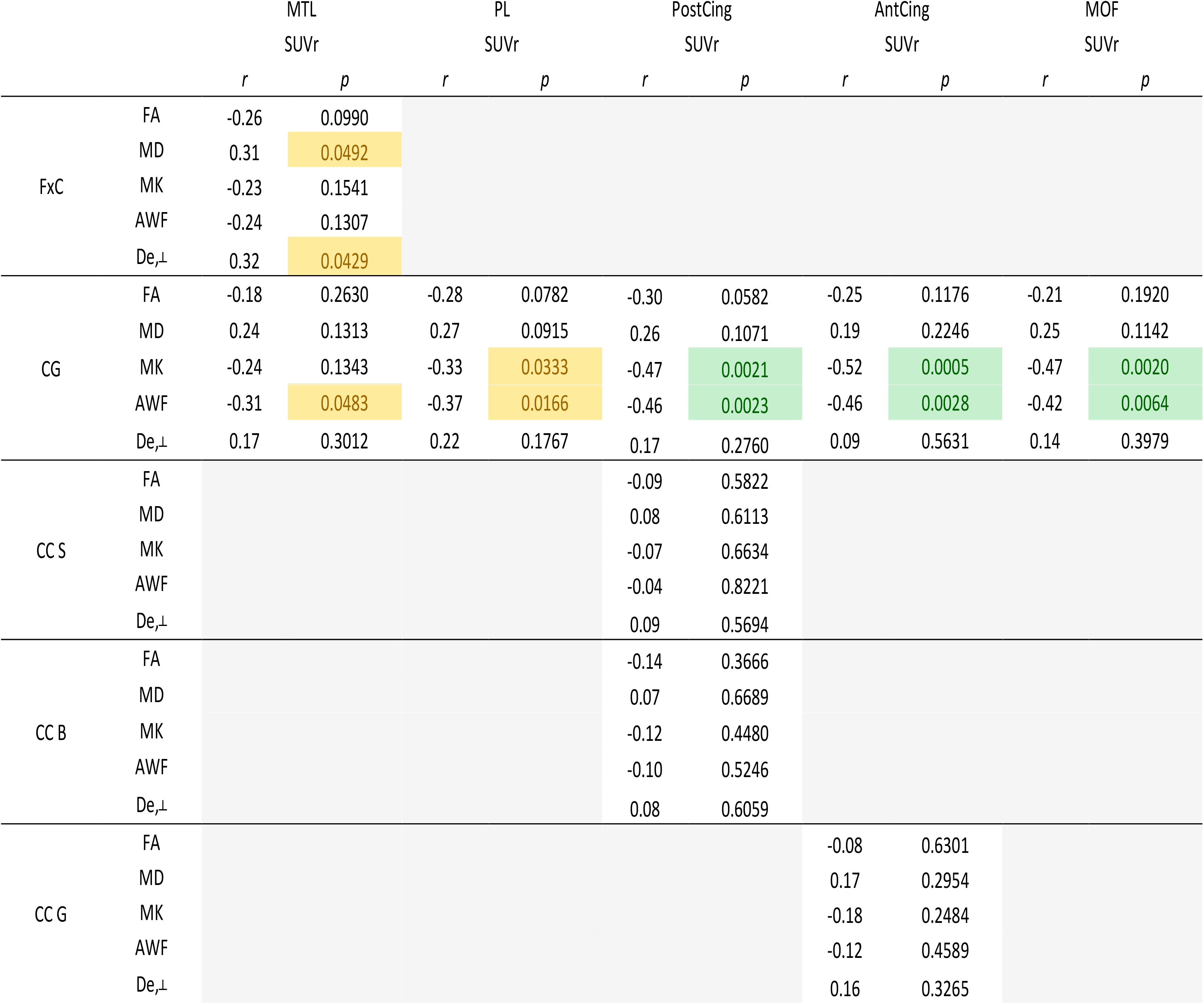
Correlations between WM diffusion metrics and SUVr of related GM regions using age as a covariate. Yellow: significant correlations (p < 0.05). Green: Still significant after Bonferroni correction (p < 0.01).

|  |  | MTL<br>SUVr |  | PL<br>SUVr |  | PostCing<br>SUVr |  | AntCing<br>SUVr |  | MOF<br>SUVr |  |
| --- | --- | --- | --- | --- | --- | --- | --- | --- | --- | --- | --- |
|  |  | *r* | *p* | *r* | *p* | *r* | *p* | *r* | *p* | *r* | *p* |
| FxC | FA | -0.26 | 0.0990 |  |  |  |  |  |  |  |  |
|  | MD | 0.31 | 0.0492 |  |  |  |  |  |  |  |  |
|  | MK | -0.23 | 0.1541 |  |  |  |  |  |  |  |  |
|  | AWF | -0.24 | 0.1307 |  |  |  |  |  |  |  |  |
|  | De,⊥ | 0.32 | 0.0429 |  |  |  |  |  |  |  |  |
| CG | FA | -0.18 | 0.2630 | -0.28 | 0.0782 | -0.30 | 0.0582 | -0.25 | 0.1176 | -0.21 | 0.1920 |
|  | MD | 0.24 | 0.1313 | 0.27 | 0.0915 | 0.26 | 0.1071 | 0.19 | 0.2246 | 0.25 | 0.1142 |
|  | MK | -0.24 | 0.1343 | -0.33 | 0.0333 | -0.47 | 0.0021 | -0.52 | 0.0005 | -0.47 | 0.0020 |
|  | AWF | -0.31 | 0.0483 | -0.37 | 0.0166 | -0.46 | 0.0023 | -0.46 | 0.0028 | -0.42 | 0.0064 |
|  | De,⊥ | 0.17 | 0.3012 | 0.22 | 0.1767 | 0.17 | 0.2760 | 0.09 | 0.5631 | 0.14 | 0.3979 |
| CC S | FA |  |  |  |  | -0.09 | 0.5822 |  |  |  |  |
|  | MD |  |  |  |  | 0.08 | 0.6113 |  |  |  |  |
|  | MK |  |  |  |  | -0.07 | 0.6634 |  |  |  |  |
|  | AWF |  |  |  |  | -0.04 | 0.8221 |  |  |  |  |
|  | De,⊥ |  |  |  |  | 0.09 | 0.5694 |  |  |  |  |
| CC B | FA |  |  |  |  | -0.14 | 0.3666 |  |  |  |  |
|  | MD |  |  |  |  | 0.07 | 0.6689 |  |  |  |  |
|  | MK |  |  |  |  | -0.12 | 0.4480 |  |  |  |  |
|  | AWF |  |  |  |  | -0.10 | 0.5246 |  |  |  |  |
|  | De,⊥ |  |  |  |  | 0.08 | 0.6059 |  |  |  |  |
| CC G | FA |  |  |  |  |  |  | -0.08 | 0.6301 |  |  |
|  | MD |  |  |  |  |  |  | 0.17 | 0.2954 |  |  |
|  | MK |  |  |  |  |  |  | -0.18 | 0.2484 |  |  |
|  | AWF |  |  |  |  |  |  | -0.12 | 0.4589 |  |  |
|  | De,⊥ |  |  |  |  |  |  | 0.16 | 0.3265 |  |  |

**Supplementary Table 3.** Correlations between WM diffusion metrics and SUVr of related GM regions, using age and GM volume as covariates. There were no significant correlations outside of the cingulum. Yellow: significant correlations (p < 0.05). Green: Still significant after Bonferroni correction (p < 0.01).

|  |  | MTL<br>SUVr |  | PL<br>SUVr |  | PostCing<br>SUVr |  | AntCing<br>SUVr |  | MOF<br>SUVr |  |
| --- | --- | --- | --- | --- | --- | --- | --- | --- | --- | --- | --- |
|  |  | *r* | *p* | *r* | *p* | *r* | *p* | *r* | *p* | *r* | *p* |
| CG | FA | -0.06 | 0.7322 | -0.18 | 0.2547 | -0.19 | 0.2361 | -0.15 | 0.3647 | -0.20 | 0.2157 |
|  | MD | 0.09 | 0.5823 | 0.11 | 0.5188 | 0.18 | 0.2763 | 0.12 | 0.4698 | 0.23 | 0.1445 |
|  | MK | -0.15 | 0.3523 | -0.25 | 0.1251 | -0.40 | 0.0108 | -0.47 | 0.0023 | -0.46 | 0.0030 |
|  | AWF | -0.20 | 0.2109 | -0.27 | 0.0960 | -0.38 | 0.0142 | -0.39 | 0.0137 | -0.42 | 0.0076 |
|  | De<sub>⊥</sub> | 0.02 | 0.9123 | 0.07 | 0.6560 | 0.09 | 0.5759 | 0.01 | 0.9438 | 0.12 | 0.4509 |

## Abbreviations

AC: Anterior Cingulate
AWF: Axonal Water Fraction
CC(G/B/S): Corpus Callosum Genu/Body/Splenium
CG: Cingulum
DTI: Diffusion Tensor Imaging
DKI: Diffusion Kurtosis Imaging
FA: Fractional Anisotropy
FxC: Fornix Crus
(a)MCI: (amnestic) Mild Cognitive Impairment
MD: Mean Diffusivity
MK: Mean Kurtosis
MOF: Medial Orbitofrontal cortex
MTL: Medial Temporal Lobe
PC: Posterior Cingulate
PH/F: Parahippocampal gyrus and fusiform
PL: Parietal Lobe
ROI: Region of Interest
SUV(r): (relative) Standardized Uptake Value
WM: White Matter
WMTI: White Matter Integrity Metrics

